# Structural sequence evolution and Computational Modeling Approaches of the Complement System in Leishmaniasis

**DOI:** 10.1101/423855

**Authors:** Prajakta Ingale, Ritika Kabra, Shailza Singh

## Abstract

The complement system acts as central arm of innate immunity that is critical to host defense as well as the development of adaptive immunity. The origins of the complement system have so far been traced, which correlates to near to the beginnings of multi-cellular animal life. Owing to the difficulty in obtaining crystal structures of GPCRs in either inactive or active state, accurate structural modeling is still highly desirable for the majority of GPCRs. In an attempt to dissect the conformational changes associated with GPCR activation, computational modeling approaches is being pursued in this paper along with the evolutionary divergence to deal with the structural variability.

## 1. Introduction

G-protein-coupled receptors (GPCRs) are the largest and most diverse family of transmembrane receptors. [1] The C5a receptor, known as complement component 5a receptor 1 (C5AR1) is a G protein-coupled receptor for C5a. [2] It is one of the major chemoattractant receptors. To date, C5aR has only been identified and cloned in mammalian species, and its evolutionary history remains ill-defined. [3,4] A relatively low homology level has been observed across the mammalian C5aR sequences; for example, human C5aR is only 65, 67, 68, and 70.8% identical with C5aR from mouse, guinea pig, dog, and rat, respectively.[5] All these C5aR molecules possess an extra-cellular N-terminal region, seven helical hydrophobic TM regions and an intracellular C-terminal domain. C5a interact with receptor protein C5a or C5aR on the surface of target cells such as macrophages, neutrophils and endothelial cells. To gain insights into the evolution and to study the divergence time between the groups of these C5aR receptor proteins, we have carried out the molecular clock analysis.

G-protein-coupled receptors (GPCRs) regulate a diverse range of intracellular signaling pathways mediated mainly by guanine nucleotide binding proteins. These pathways form an intricate network involved in critical biological tasks but not limited to homeostasis and transcriptional factors etc. Given the central role of GPCRs in the overall biology of the cell, they undoubtedly represent the most relevant group of therapeutic targets. As stated the cross-talk network is an intricate means of regulation, synergizing weak stimuli and reigning in collateral damage. Leishmania is already well known to exploit these interactions with a multitude of evasion tactics, working to reduce its detection and prolong its survival. Most of the described mechanisms show the coupling of inhibitory receptors that dampen cellular immunity or misguide the attack. One well described method appropriates the complement system, which is among the first of the parasite host interactions. Seemingly in conflict with the definition of ‘evasion’ Leishmania of all species, potently and selectively activates complement. This selectivity is found between the various stages of the parasite, whereby promastigotes and amastigotes can differentially target the human complement receptors CR1 and CR3 to suit their distinct living conditions. This cunning use of cross talk increases phagocytosis through opsonized complement receptor mediated uptake and decrease the intensity of the oxidative burst within the phagosome. It is so pivotal to pathogenesis that genetic ablation of the complement receptor C5aR renders mice resistant to infection with *L. major*. [6]

Owing to the difficulty in obtaining crystal structures of GPCRs in either inactive or active state, accurate structural modeling is still highly desirable for the majority of GPCRs. Over the years, structural modeling of GPCRs has mostly been obtained through homology modeling using available crystal structures as suitable templates but various *ab initio* approaches have also been tested. Computational methods that have been applied in the course of these years to study the dynamical properties of GPCRs in an attempt to dissect the conformational changes associated with their activation, is being pursued in this paper along with the molecular clock analysis.

## 2. Materials and methods

### 2.1. Data Collection

Total 22 coding sequences of C5AR1 genes (of different vertebrates) were retrieved form gene database of NCBI in .fasta file format. These .fasta files merge together to form multifasta file. The accession no. of different organism’s C5AR1 included for the analysis is given as below;

| Sr. No. | Accession No. | Scientific Name | Common Name |
| --- | --- | --- | --- |
| 1. | GI\|291167767\| | *Mus musculus* | House mouse |
| 2. | GI\|112293258\| | *Homo sapiens* | Human |
| 3. | GI\|847158952\| | *Xenopus tropicalis* | Frog |
| 4. | GI\|297277451\| | *Macaca mulatta* | Monkey |
| 5. | GI\|694975876\| | *Pan troglodytes* | chimpanzee |
| 6. | GI\|545175925\| | *Equus caballus* | Horse |
| 7. | GI\|16758417\| | *Rattus norvegicus* | Rat |
| 8. | GI\|166999226\| | *Bos taurus* | Cattle |
| 9. | GI\|50979237\| | *Canis lupus familiaris* | domestic dog |
| 10. | GI\|346644763\| | *Sus scrofa* | wild boar |
| 11. | GI\|290543381\| | *Cavia porcellus* | Guinea pig |
| 12. | GI\|744599290\| | *Camelus dromedarius* | Camel |
| 13. | GI\|548505144\| | *Capra hircus* | Goat |
| 14. | GI\|591293794\| | *Panthera tigris altaica* | Tiger |
| 15. | GI\|564240887\| | *Alligator mississippiensis* | Alligator |
| 16. | GI\|587013626\| | *Felis catus* | Cat |
| 17. | GI\|655892848\| | *Oryctolagus cuniculus* | Rabbit |
| 18. | GI\|752416049\| | *Ailuropoda melanoleuca* | Giant panda |
| 19. | GI\|688598194\| | *Danio rerio* | Zebrafish |
| 20. | GI\|602652885\| | *Python bivittatus* | Python |
| 21. | GI\|675726191\| | *Pan paniscus* | Bonobo |
| 22. | GI\|426389371\| | *Gorilla* | Gorilla |

### 2.2 Sequence alignment and phylogenetic reconstruction

Multiple sequence alignment was performed using standalone clustalW2 program [7] and the .aln file generated was converted to .nexus format to create Bayesian inference phylogeny and molecular clock using MrBayes (Version 3.2.2 ×64). [8, 9] The final trees generated by the MrBayes were visualized and edited using Fig tree software (vs 1.4.2). [10]

#### 2.2.1. Bayesian inference phylogeny and Relaxed Clock Model

To estimate Bayesian inference phylogeny, GTR + I +G model was used and Markov chain Monte Carlo (MCMC) simulation was run for 30,000 generations, until the standard deviation of split frequencies were below 0.01. Parameters and corresponding tree were summarized after discarding the initial 25% of each chain as burnin. Information regarding the out-groups was obtained using the resulting tree generated by Bayesian inference method. Relaxed molecular clock was constructed using the same dataset by running 4,00,000 generations of MCMC simulation. The model used for constructing the molecular clock was the independent gamma rates model (IGR model).

#### 2.2.2 Node dating of relaxed molecular clock

We have calibrated the relaxed clock model based on the node calibration (node dating). Based on the default setting e. g, the clock rate is fixed to 1.0, means the age of the nodes in the tree will be measured in the number of expected substitution per site per million years, we assume that the rate is approximately 0. 01 ± 0.005 substitutions per site per million year. For this we have use 0.02 as the mean and 0.005 as the standard deviation. Since we give these values using millions of years as the unit, the resulting tree will be calibrated in millions of years.

### 2.3 Homology Modeling and Molecular Dynamics Study

The primary amino acid sequence of human C5AR1 was downloaded from NCBI gene sequence database (NP_001727). Visualization and presentation of model structure was performed in Pymol (PyMOL Molecular Graphics System, Version 1.7, Schrödinger, and LLC). Ab-initio modeling was carried out using online I-TASSER server. [11] Loop refinement and quality assessment was performed using MODELLER and also other online server like SAVES, PDBsum [12] RAMPAGE etc. [13, 14] and finally the energy minimization and atomistic simulation study was performed using GROMACS [15].

### 2.4 Molecular dynamics simulation of C5aR1

MD simulation was performed using OPLS-AA (all-atom) force field and the SPC (Simple Point Charge) water model. The protein was solvated with water in periodic cubic box of system size 77.74 Å × 51.32 Å × 90.78 Å and the minimal distance of 1.0 nm (10 Å) was applied between the edge of the box and the protein. First the energy minimization was carried out, which has taken 2329 steps of steepest descent to get converged. After minimization potential energy noted was −3.8550902e+06. The temperature and pressure was controlled with the v-rescale and Berendsen weak coupling algorithms. Finally the production MD was performed for 12ns time steps until the model gets stabilized. After completion of the simulation steps, the trajectory of the protein, energy and RMSD (root-mean-square-deviation), RMSF (root-mean-square-fluctuation) were analyzed.

#### C5aR1 Lipid Bilayer (POPC) Membrane MD Simulations

After successful MD simulation minimum energy conformation of C5aR1 structure was obtained from trajectory. C5aR1 belongs to the family of GPCRs; their 7 transmembrane helical domains were identified visually by analyzing the structure. For the Lipid bilayer membrane insertion POPC (1-palmitoyl-2-oleoyl-sn-glycero-3-phosphocholine) [17] was selected due to its versatile vesicle formation ability. Lipid bilayer was inserted using Desmond (DE Shaw Research, USA) software System Builder module. Total 182 POPC molecules were inserted to cover the entire 7 transmembrane helices of C5aR1. The area per lipid of POPC was around 68.2 Å□^2^. After insertion of POPC bilayer TIP3P [18] water model was used for solvation. Other parameters used were kept constant which were mentioned in earlier MD simulation section. A total of 5ns of production MD run was performed with 2fs of time step and snapshots were saved for every 2ps. At the end of the MD simulation POPC area per lipid was close to experimentally determined areas (68.3 ± 1.5 Å□^2^), thus indicating that this lipid bilayer MD simulation is successful.

## 3. Result and Discussion

### 3.1 Bayesian inference phylogeny and Relaxed clock model

**Figure 1:**
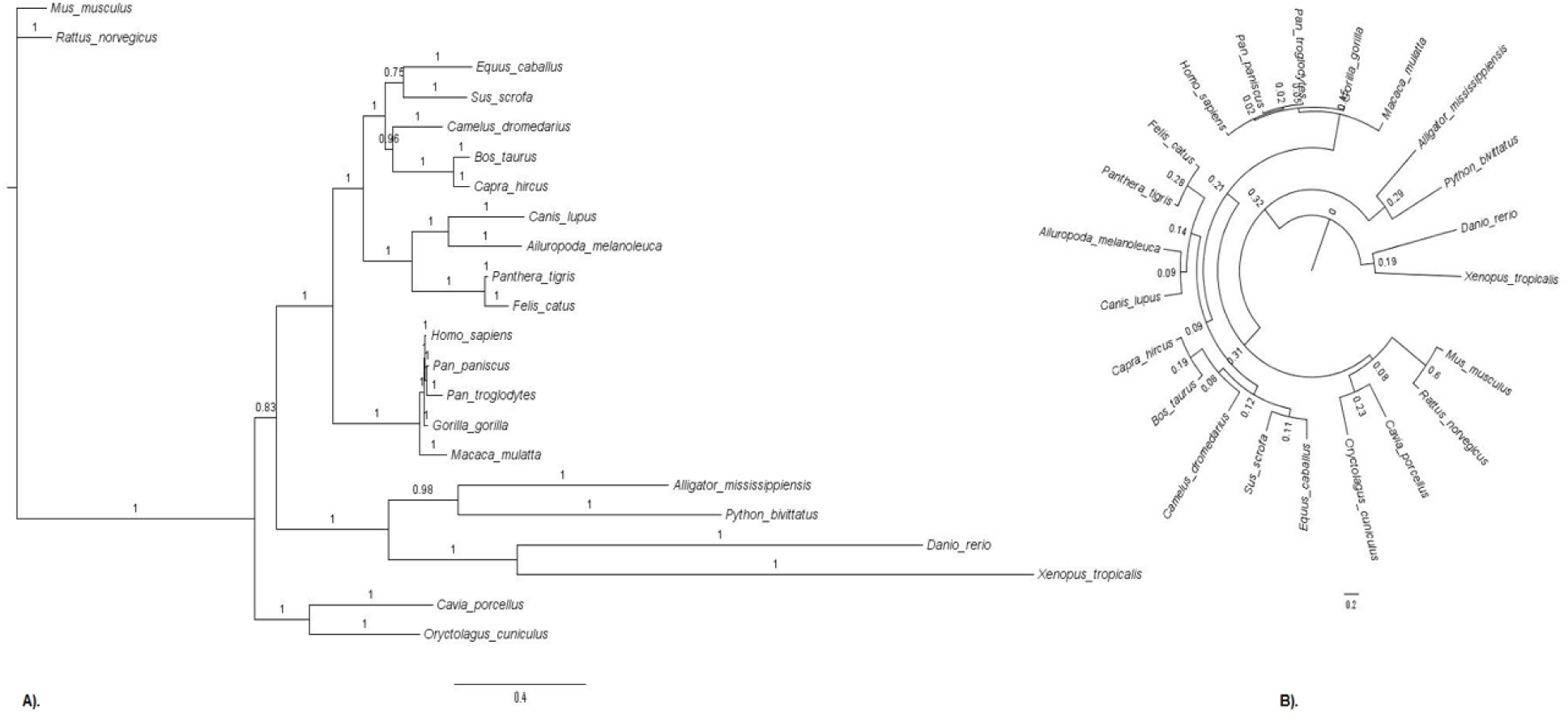
**A).** Phylogram based on Bayesian analysis of 22 taxa of C5AR1 genes of different organisms. The horizontal lines are branches and represent evolutionary lineages changing over time. The longer the branch in the horizontal dimension, the larger the amount of change. The bar at the bottom of the figure provides a scale for this. In this case the line segment with the number ‘0.4’ shows the length of branch that represents an amount genetic change of 0.4. Support values on branches are Bayesian posterior probabilities (BPPs). A high value means that there is strong evidence that the sequences to the right of the node cluster together to the exclusion of any other. The species *Ratus norvegicus* and *Mus musculus* were included as out-group. **B).** Relax clock model based on the Bayesian tree. The support value at each node indicates the branch length that is proportional to time. The scale bar at the bottom of the figure represents the amount of change that is 0.2. Here we can see that relaxed clock model generated was accelerating with respect to the out-group.

### 3.2 Node Age Chronogram

**Figure 2:**
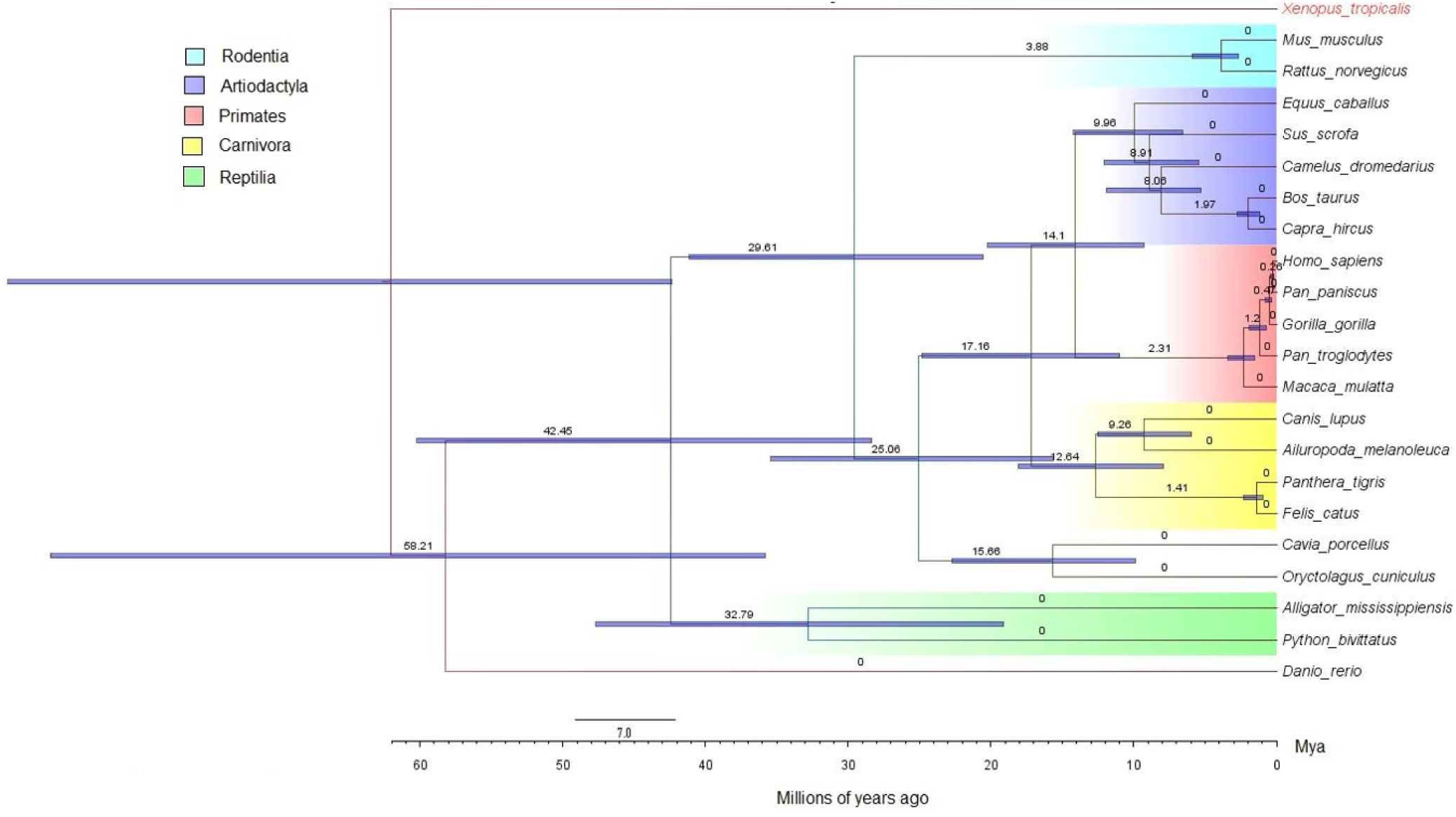
Chronogram of Relaxed clock model based on node dating (ND). Here the node labels indicate the median age (mya) and gray node bars shows the 95% HDP of the estimated node age. Out-group taxa used to root the tree include *Xenopus tropicalis*. The clades were marked with different shades of colors corresponding to the distribution of the taxa according to their order. The divergence time is indicated in Millions of years (Mya) below the figure. Scale bar represents the number of substitutions per site.

### 3.3 Molecular Modeling of C5aR1

In order to carry out homology modeling using MODELLER of C5aR1, best template selection was performed through PSI BLAST of the target protein against the PDB database (http://www.rcsb.org/pdb/home/home.do). The significant hits with ≥30% sequence identity, was selected as templates for the target protein. The blast result shows that 4DKL allow relatively high (30%) sequence similarity with C5aR1. After generating the homology based model using MODELLER it was observed that the N-terminus region of C5aR1 was not able to span over the entire 4DKL, as a result the secondary structure of initial 34 residue of N-terminus was not obtained. So *ab*-*initio* modeling using I-TASSER (Iterative Threading Assembly Refinement) server was performed.

Three criteria (C-Score, TM score and RMSD values) were employed to compare the quality of predicted models to choose the best one. Model 1 was predicted to have a C-Score −0.42 which was comparatively better than the other homology model. The estimated computational TM score 0.66±0.13 and RMSD 7.4±4.3 of model 1 was found to be better than the values obtained for other model.

I-TASSER provides best possible 5 models; we can choose any one out of these 5 structures. This server provides the TM score and RMSD for only first structure although the other structures can also be considered if it fulfills the other criteria. Here, the below table provides the 5 structures generated by I-TASSER and their different parameter values.

Following are the results obtained by I-TASSER server:

**Table 1:**
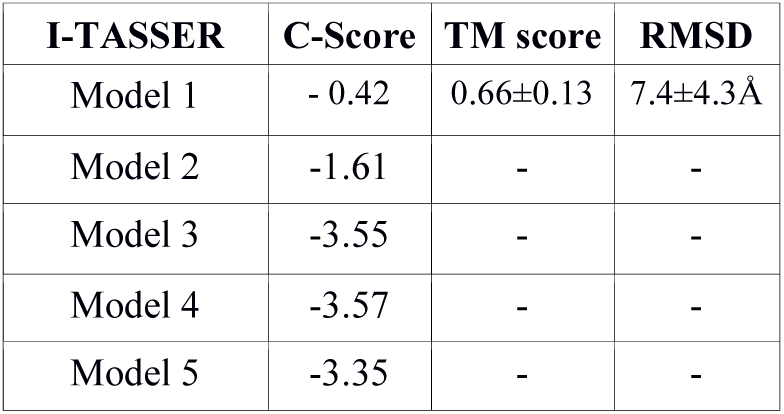
Three criteria (C-Score, TM score and RMSD values) of I-TASSER After analyzing I-TASSER parameters, these structures were further assessed through Ramachandran plot using RAMPAGE, dope score and GA341 score using MODELLER. The result obtained after the analysis is shown as below:

| **I-TASSER** | **C-Score** | **TM score** | **RMSD** |
| --- | --- | --- | --- |
| Model 1 | - 0.42 | 0.66±0.13 | 7.4±4.3Å |
| Model 2 | -1.61 | - | - |
| Model 3 | -3.55 | - | - |
| Model 4 | -3.57 | - | - |
| Model 5 | -3.35 | - | - |

**Table 2:** Result of RAMPAGE and MODELLER modules

| **I-TASSER**<br>**Models** | **RAMPAGE** |  |  | **Dope score** | **GA341** |
| --- | --- | --- | --- | --- | --- |
|  | **Favored** | **Allowed** | **Outliers** |  |  |
| Model 1 | 292 | 40 | 16 | - 45573.589844 | 1.000000 |
| Model 2 | 289 | 43 | 16 | - 42572.429688 | 1.000000 |
| Model 3 | 267 | 59 | 22 | - 44035.679088 | 1.000000 |
| Model 4 | 288 | 38 | 22 | - 43878.457031 | 1.000000 |
| Model 5 | 297 | 36 | 15 | - 44077.480469 | 1.000000 |

Based on above analysis, it can be said that the value of dope score and GA341 score was within the limit but the arrangement of residues based on Ramchandran plot required loop refinement. Visualization of the structure was performed using Pymol. [16]. Model 4 showed the structural correlation with the predefined structure of model C5aR1 as it contains 7 helical domains, 3 intra and 3 extra cellular loops **(Figure 3)**. Initial refinement of 3D model generated was carried out with the help of loop refinement protocol of MODELLER. For this script loop.py was run in order to move 22 residues from the outliers to allowed region. The final structure obtained was again analyzed for its quality assessment.

**Figure 3:**
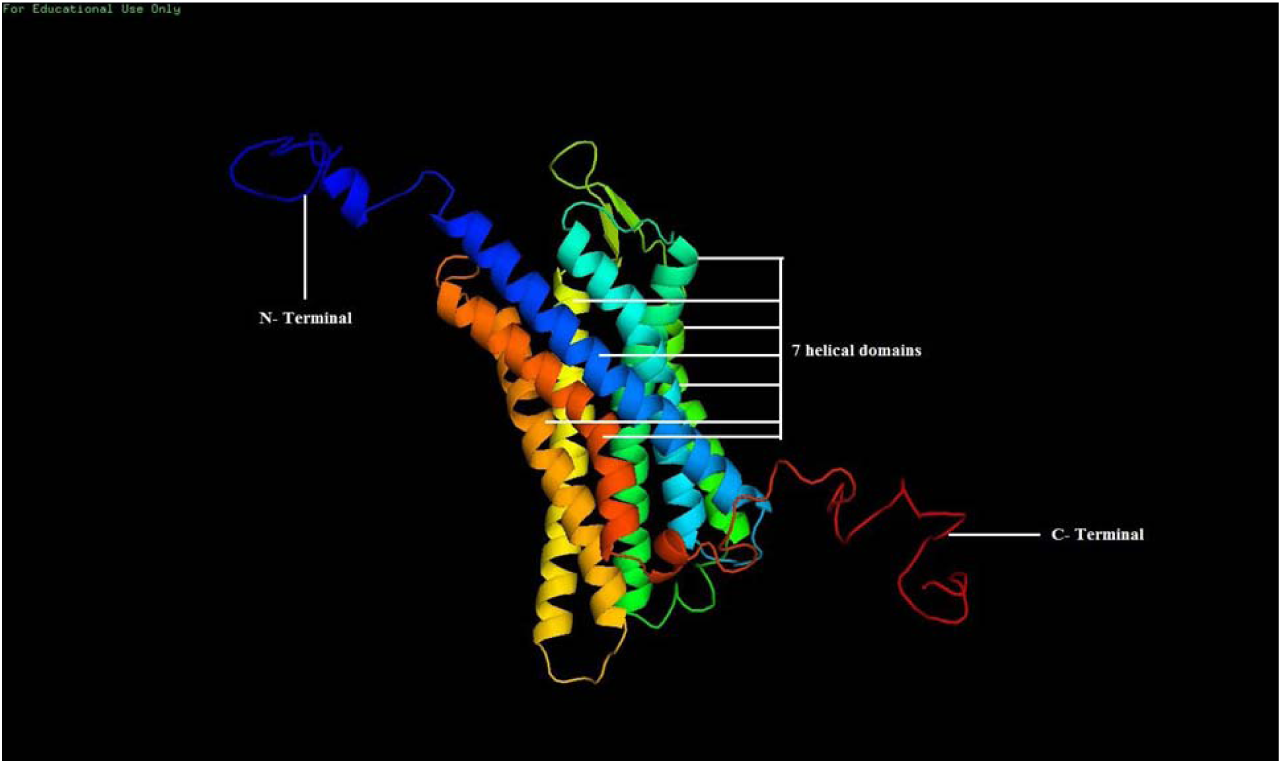
Final model (model 4) of C5aR1.pdb visualized through pymol. Here initial blue part represents the free N-terminal region where as red part shows the free C-terminal region, the middle region has 7 helical domains, showing resemblance with the predetermined theoretical model.

**Table 3:** Result of RAMPAGE and MODELLER modules.

| **I-TASSER**<br>**Final model**<br>**4.pdb** | **RAMPAGE** |  |  |
| --- | --- | --- | --- |
|  | **Favored** | **Allowed** | **Outliers** |
|  | 322 | 26 | 0 |
| Dope score | -42919.250 |  |  |
| GA341 | 1.00 |  |  |

After the loop refinement and quality assessment using MODELLER, the final model was verified by Structural Analysis and Verification Server (SAVES) to evaluate its stereo-chemical quality. Ramachandran plot computed by PROCHECK module and other features of the structures were analyzed by different structure prediction servers. Following **(Figure 4)** are the results of final model using different programs.

**Figure 4:**
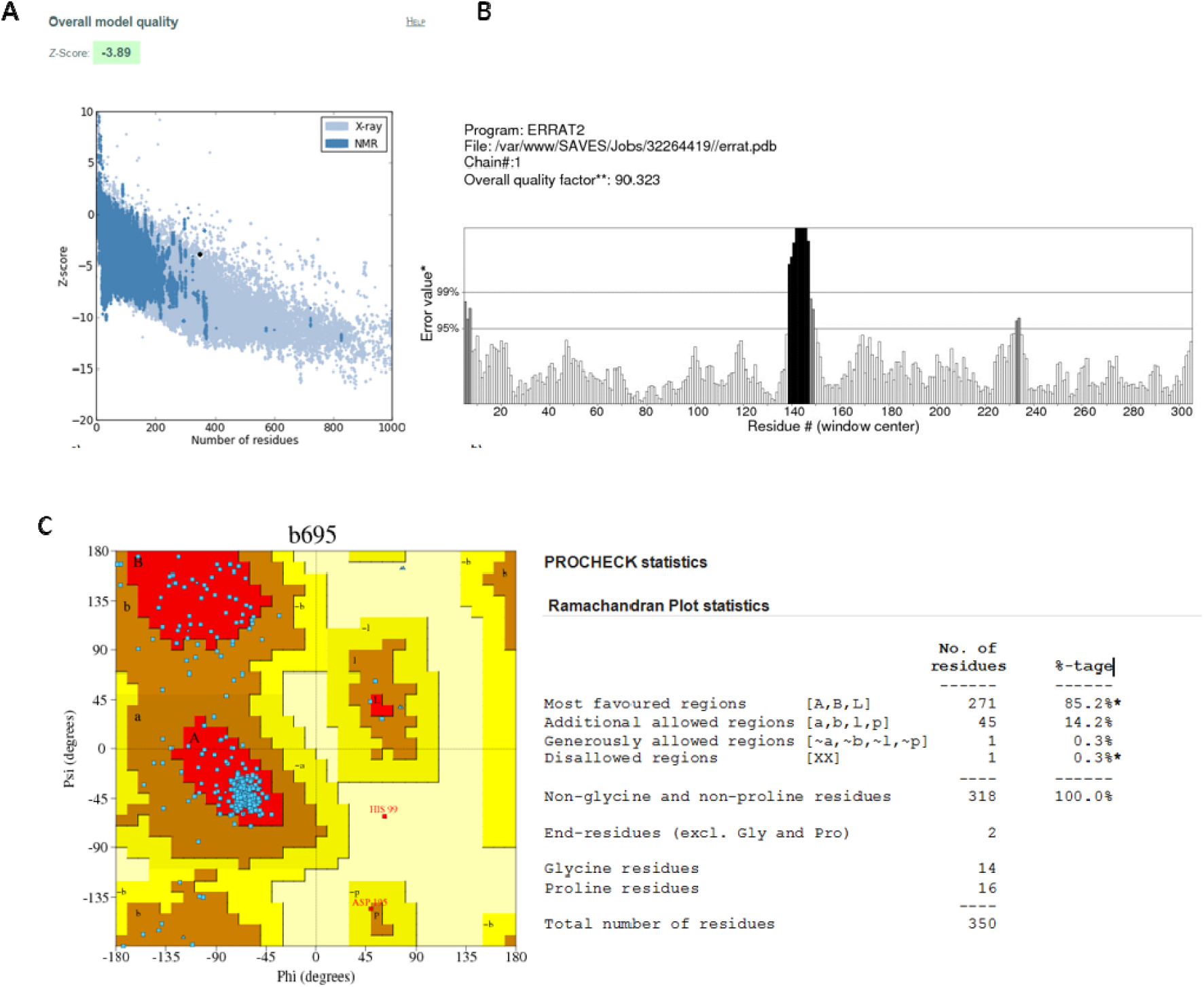
**a).** ProSAweb result shows the overall quality of final structure. **(b).** Errat result by SAVES (Structural Analysis and Validation Server) results. **c).** Ramachandran plot by PROCHECK.

Evaluation of model-4 structural quality with ProSA-web (**Figure 4a**) revealed its ProSA Z-score value −3.89, falls in the range of native conformations computed using X-ray crystallography method represented as encircled as large black dot. Errat result shows the overall quality of the structure is 90.323 (**Figure 4b**), which is very good experimentally and computationally. Ramachandran plot computed by PROCHECK module **(Figure 4c)** showed only 0.3% of residues exist in disallowed regions confirming the quality of 0.3% residues predicted to be highly significant.

### 3.4 Molecular dynamics simulation of C5aR1

The resulting plots of each stage of MD simulation are represented as below:

From the above figures **(Figure 5A)** The Energy graph of Potential demonstrating steady convergence of Epot and from the temperature plot **(Figure 5B)**, it is clear that the temperature of the system quickly reaches the target value (300K), and remains stable over the remainder of equilibration. From the Pressure plot **(Figure 5C)**, pressure value fluctuates widely over the course of 100-ps equilibration phase, but this behavior is not unexpected and from the density plots **(Figure 5D)**, we can clearly see that density values are very stable over time, indicating that the system is well-equilibrated now with respect to pressure and density.

**Figure 5:**
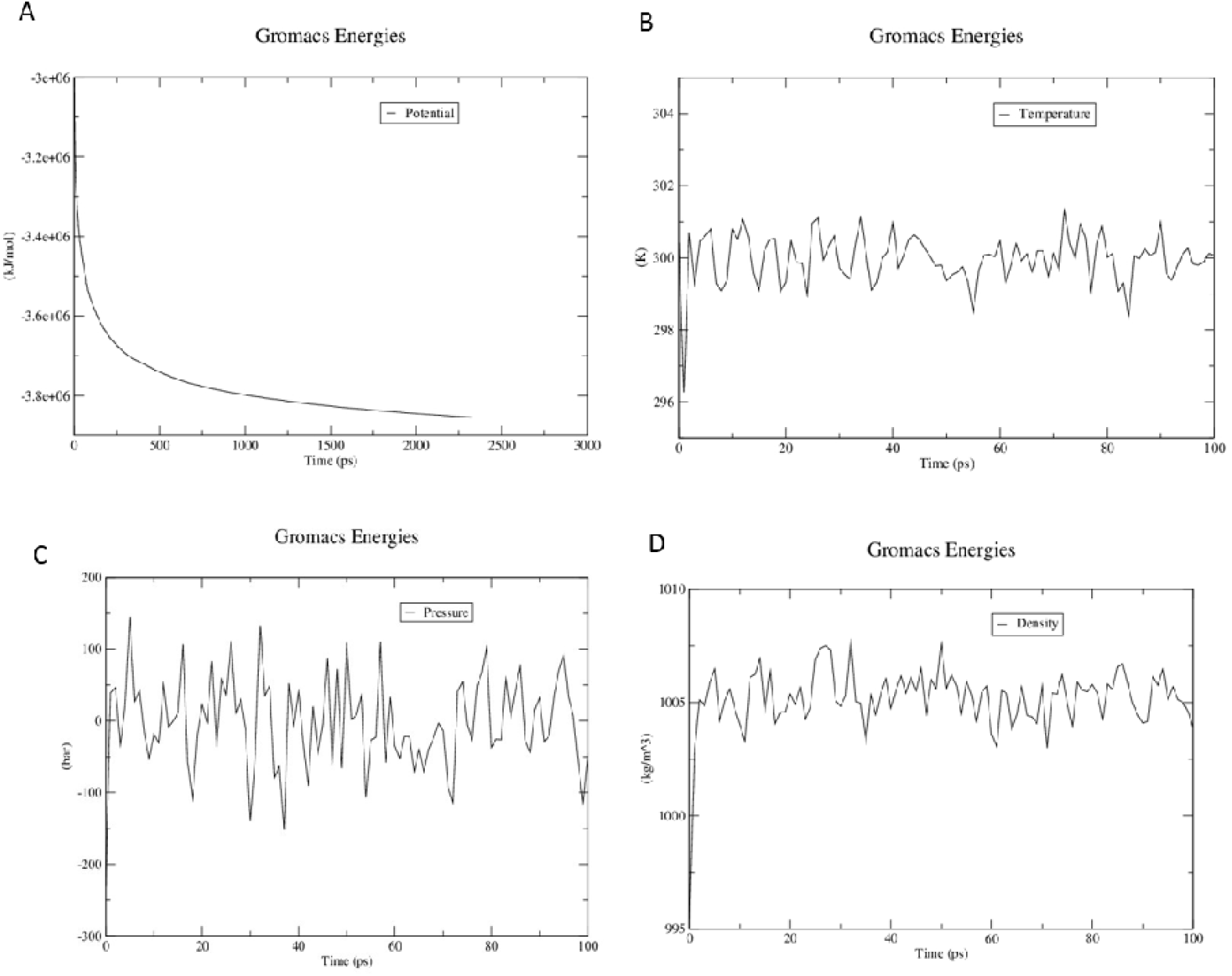
Parameters during MD simulation A) Energy minimization B) Temperature C) Pressure D) Density plot.

Here, above the first figure **(Figure 6A)** shows all-atom RMSD of the modeled system against the time scale. After 6 ns time the system starts stabilizing, it can be visualized that after 6ns the deviation was found to occur within a range of 0.7-0.8□nm, means within 1□Å, it can be regarded as stable. The second plot **(Figure 6B)** shows the root mean square fluctuation (RMSF) for each atom. It also shows flexibility for each atom but at some point (i. e. atom no. 3000), it gets fluctuated.

**Figure 6:**
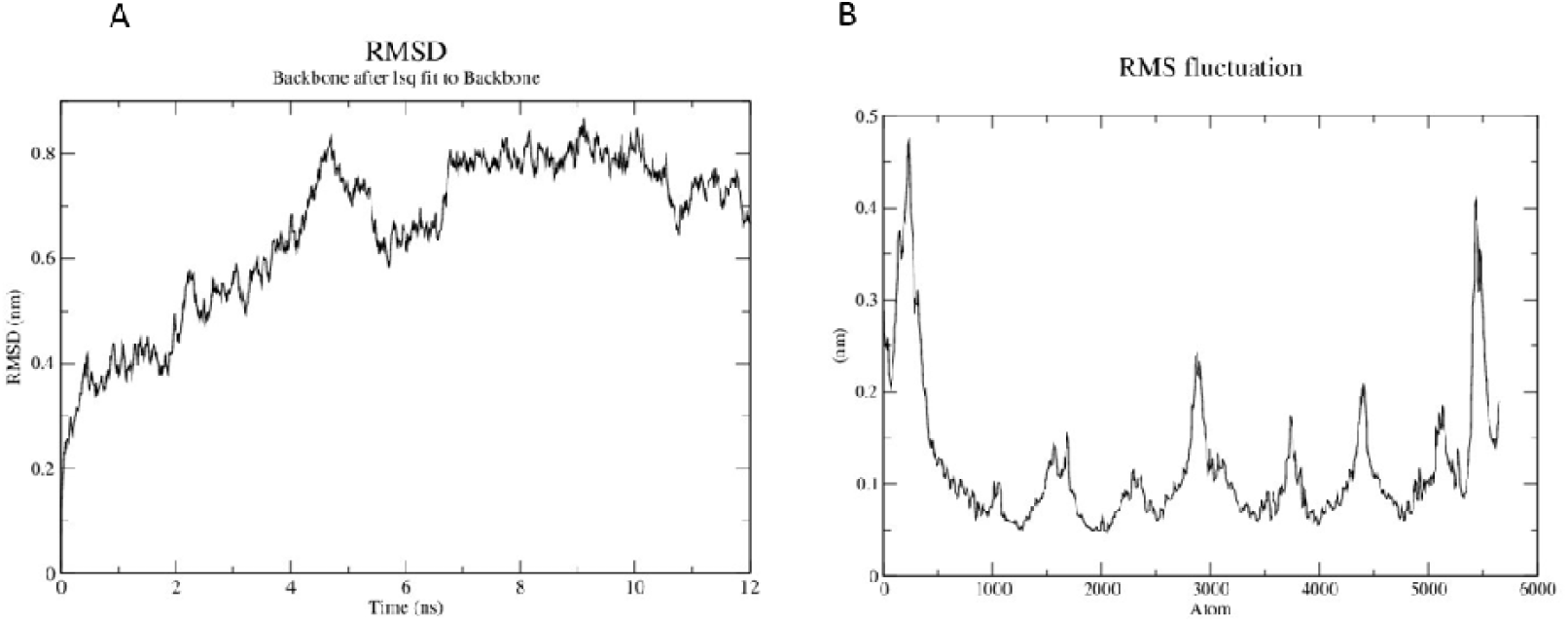
A) RMSD plot B) RMSF plot.

**Fig. 7.**
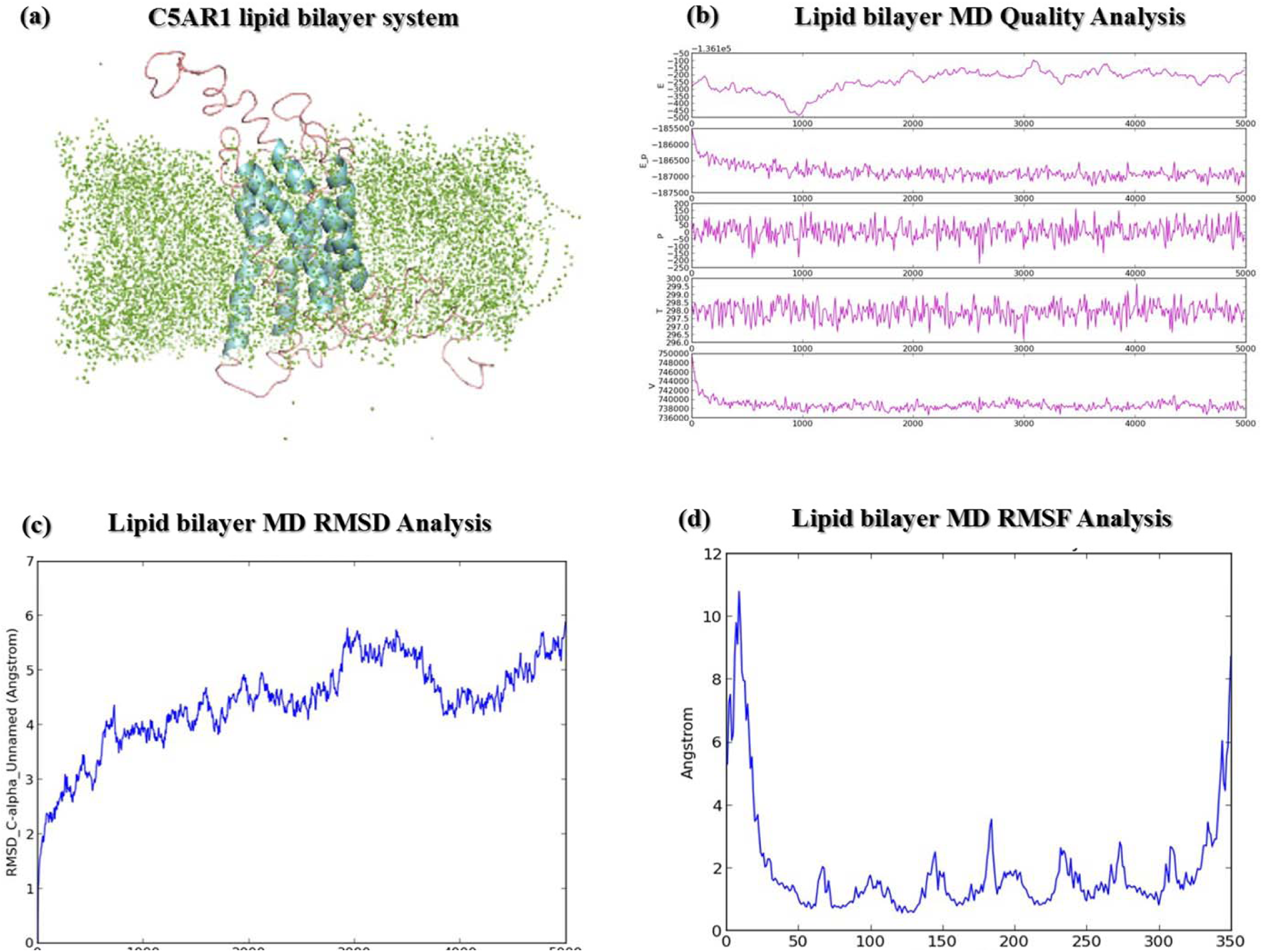
a) C5aR1 Lipid Bilayer system (b) Quality plot generated for C5aR1(c) Lipid Bilayer MD RMSD analysis (d) Lipid bilayer MD RMSFanalysis.

## 4 Discussion

In this study, we aimed to provide deeper insight into the species diversity of C5AR1 receptor genes of different groups. This study helped us to clarify some aspects of the evolutionary history of this interesting group. Based on the Bayesian inference phylogeny (**Figure 1A**), the Bayesian posterior probabilities (BPPs) values is almost 1 at every node, indicated that clade generated were the closest relatives among all the species used for the phylogenetic reconstruction. From the relaxed molecular clock (**Figure 1B**), it can be said that the clock is accelerating with respect to the out-group. The clades are highlighted with the different color in the final chronogram which confirmed that *Rodentia*, *Artiodactyla*, *Primates*, *Carnivora* and *Reptilia* form different clades according to their orders. The divergence times and the mean age estimates of most nodes were largely consistent. The phylogram gives the median age estimates for all nodes in the tree, which is million years. On the basis of chronogram (**Figure 2**), we can see that the oldest spilt in the tree is at around 63 Mya (million years ago). The molecular dating results indicate that clade formed for *Rodentia* (highlighted with blue) diverged approximately 29.61 mya (95% Highest Posterior Density) from the other groups. It may be stated that within the *Rodentia* clade, the taxa *Mus musculus* and *Rattus norvegicus* diverged around 3.88 mya from each other, while clads highlighted with purple (*Artiodactyla*) and pink (*Primates*) diverged at 14.1 mya. The taxa within the group *Carnivora* and *Reptilia* diverged at approximately 12.64 and 32.79 mya respectively. Divergence among the individuals of six species (Homo *sapiens*, *Pan paniscus*, *Gorilla*, *Pan troglodytes* and *Maccaca mulatta*) were between 0.26 mya, 1.2 mya and 2.3mya respectively. The developed model of C5aR1 shows overall good structural quality and was confirmed using several different validation tools. It can act as a good chemo attractant target as C5a interacts with receptor protein C5a1 or C5aR on the surface of target cells such as macrophages, neutrophils and endothelial cell. In Leishmania infection, the amastigote form of the pathogen is found present in human macrophages. So by targeting this C5AR receptor which is one of the most potent inflammatory chemoattractant and has been implicated in pathogenesis of numerous inflammatory diseases, we can further stop the cell division of amastigote form of Leishmania. Our study provides the molecular model and its *in*-*silico* structural stability but still further study needs to be done in order to understand the biochemical nature of the molecule. C5aR1 has raised the intriguing possibility of the use of this receptor having possible role in inflammation.

## 5. Acknowledgement

The authors would like to thank Department of Biotechnology, Government of India for intramural fund.

